# Excess glutamate release triggers subunit-specific homeostatic receptor scaling

**DOI:** 10.1101/2022.05.03.490370

**Authors:** Pragya Goel, Yifu Han, Nancy Tran, Samantha Nishimura, Sarah Perry, Manisha Sanjani, Dion Dickman

## Abstract

Ionotropic glutamate receptors (GluRs) are targets for modulation in Hebbian and homeostatic synaptic plasticity and are remodeled by development, experience, and disease. Although much is known about activity-dependent mechanisms that regulate GluR composition and abundance, the role of glutamate itself in these processes is unclear. To determine how glutamate sculpts GluR receptive fields, we have manipulated synaptically released glutamate and generated precise CRISPR mutations in the two postsynaptic GluR subtypes at the *Drosophila* neuromuscular junction, GluRA and GluRB. We first demonstrate that GluRA and GluRB compete to establish postsynaptic receptive fields, and that proper GluR abundance and localization can be orchestrated in the absence of any synaptic glutamate release. However, excess glutamate release adaptively tunes postsynaptic GluR abundance, echoing GluR receptor scaling observed in mammalian systems. Unexpectedly, when GluRA vs GluRB competition is eliminated, excess glutamate homeostatically regulates GluRA abundance, while GluRB abundance is now insensitive to glutamate modulation. Finally, Ca^2+^ impermeable GluRA receptors are no longer sensitive to homeostatic regulation by glutamate. Thus, excess glutamate, GluR competition, and Ca^2+^ signaling collaborate to selectively target GluR subtypes for homeostatic regulation at postsynaptic compartments.

## INTRODUCTION

Ionotropic GluRs are dynamically regulated at postsynaptic densities during development, plasticity, aging, and disease. A variety of intracellular signaling systems in the postsynaptic compartment coordinate the establishment, maintenance, and remodeling of GluR abundance and composition during development, plasticity, and disease (Diering and Huganir, 2018; Herring and Nicoll, 2016). Another layer of regulation involves competition between GluR subtypes, where, for example, Ca^2+^ permeable and impermeable AMPA receptor subtypes are selectively modulated during plasticity (Herring and Nicoll, 2016; Park et al., 2018), and alterations in synaptic activity can drive both Hebbian and homeostatic remodeling of GluRs (Chowdhury and Hell, 2018; Diering and Huganir, 2018; Li et al., 2019; Pozo and Goda, 2010; Turrigiano, 2008). The role of the neurotransmitter glutamate itself in regulating postsynaptic GluRs is somewhat counterintuitive: While glutamate is dispensable for the development of GluR fields (Sando et al., 2017; Sigler et al., 2017), extracellular glutamate alone is capable of provoking GluR assembly *de novo* (Kwon and Sabatini, 2011). To what extent levels of synaptic glutamate itself influences GluR composition and abundance has not been clearly defined.

The glutamatergic *Drosophila* neuromuscular junction (NMJ) is an attractive system to interrogate the role of synaptic glutamate in establishing and adaptively modulating postsynaptic GluR fields. At this synapse, the postsynaptic GluRs are classified as kainate-type GluRs (KARs; (Li et al., 2016)), tetramers composed of three essential subunits (GluRIIC, GluRIID, and GluRIIE) and one of two alternative subunits, GluRIIA or GluRIIB ((Marrus et al., 2004; Qin et al., 2005); Fig. 1A). Here, we abbreviate these distinct KAR subtypes as “GluRA” and “GluRB” to define receptors containing either the GluRIIA or GluRIIB subunit. Studies *in vivo* and in heterologous systems have shown that the majority of postsynaptic Ca^2+^ influx and depolarizing currents are driven by GluRA receptors, while GluRB passes much less current due to rapid desensitization (Diantonio et al., 1999; Han et al., 2015). In addition, the recent development of a botulinum neurotoxin that blocks neurotransmitter release in *Drosophila* now enables the silencing of synaptic glutamate release (Han et al., 2022), while excess glutamate can be released from individual synaptic vesicles by overexpression of the *vesicular glutamate transporter* in *Drosophila* motor neurons (Daniels et al., 2004; Gaviño et al., 2015; Li et al., 2018, 2021). However, to what extent changes in synaptic glutamate levels are capable of adaptively modulating GluRs at the fly NMJ is not known.

**Figure 1:**
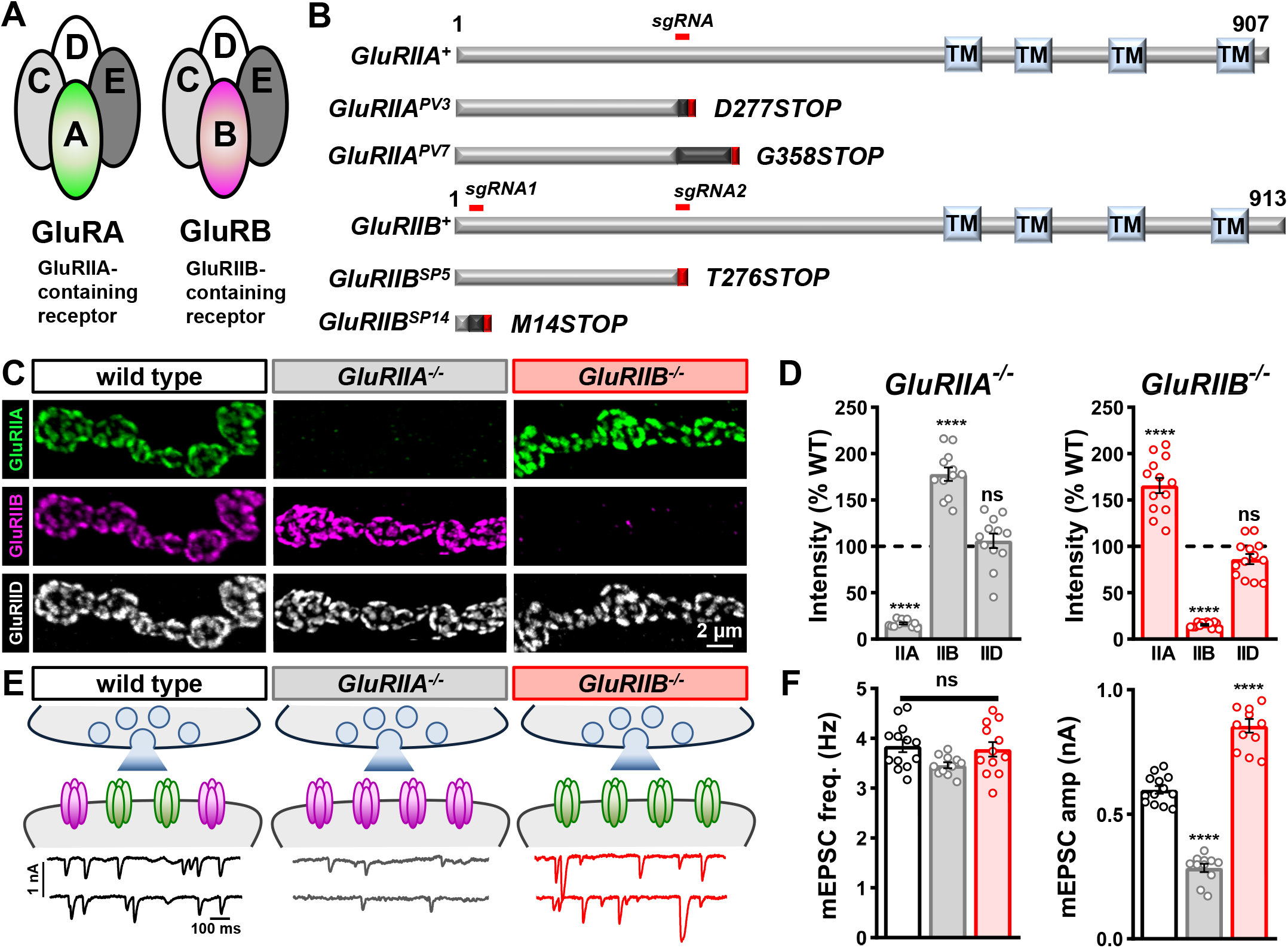
GluRIIA- and GluRIIB-containing receptor subtypes compete to establish postsynaptic glutamate receptor fields. **(A)** Schematic illustrating the subunit composition of GluRIIA- and GluRIIB-containing postsynaptic GluR subtypes at the *Drosophila* NMJ, referred to here as GluRA and GluRB receptors. **(B)** Domain structure of GluRIIA and GluRIIB subunits with the region targeted by the single guide (sg)RNA used to generate the CRISPR mutant alleles and their predicted protein products indicated below. **(C)** Representative images of muscle 4 NMJs in wild type (*w^1118^*), *GluRIIA* mutants (*w;GluRIIA^PV3^*), and *GluRIIB* mutants (*w;GluRIIB^SP5^*) immunostained with antibodies against three postsynaptic GluR subunits (GluRIIA, GluRIIB, and GluRIID). **(D)** Quantification of mean GluR fluorescence intensity normalized to wild type values confirms GluRIIA subunits are not detected at NMJs of *GluRIIA^pv3^* mutants, while GluRIIB levels are significantly increased. An inverse change is found in *GluRIIB^sp5^* mutants. Similar results are observed in *GluRIIA^pv7^* and *GluRIIB^sp14^* alleles. Overall levels of the common GluRIID subunit do not significantly change in either mutant. **(E)** Schematic summarizing the data in (C,D). Representative electrophysiological traces of mEPSC events in the indicated genotypes are shown below. **(F)** Quantification of mEPSC frequency and amplitude in the indicated genotypes. Note that mEPSC amplitude is significantly reduced in *GluRIIA^PV3^* and increased in *GluRIIB^SP5^* mutants, consistent with the staining results.

We have generated the first null mutations that specifically ablate *GluRIIA* and *GluRIIB* receptor subunits using CRISPR/Cas9 gene editing. These mutants have provided an opportunity to probe how GluR fields are established during development, how they respond to synaptically released glutamate, and to define how competition between GluRA and GluRB influences the postsynaptic response to glutamate on GluR plasticity. These studies reveal a hierarchy of control through competition between GluR subtypes and selective, homeostatic regulation by synaptic glutamate that requires Ca^2+^ permeability through GluRA receptor subtypes.

## RESULTS

### *GluRIIA* and *GluRIIB* mutants reveal competition between receptor subtypes

To understand how postsynaptic receptive fields are established at the *Drosophila* NMJ, we generated targeted genetic mutations in the two distinctive GluR subunits, *GluRIIA* and *GluRIIB* (Fig. 1A,B). Although a mutant allele of *GluRIIA* was generated over 20 years ago using imprecise transposon excision (*GluRIIA^SP16^*; (Petersen et al., 1997)), this lesion did not cleanly disrupt *GluRIIA* only, with expression of a neighboring gene, *oscillin*, also disrupted (Chen and Dickman, 2017). Specific mutations in *GluRIIB* only have not been reported. We used single guide RNAs (sgRNAs) targeting early exons of the *GluRIIA* or *GluRIIB* coding regions combined with Cas9-mediated mutagenesis to generate a series of specific mutations in either subunit; two independent null mutations in *GluRIIA* and *GluRIIB* were chosen for further analysis (see methods; Fig. 1B). To validate these alleles, we co-immunostained the larval NMJ with antibodies against GluRIIA, GluRIIB, and the common essential subunit GluRIID. As expected, GluRIIA was absent in the *GluRIIA* mutant, GluRIIB was absent in the *GluRIIB* mutant, and the common GluRIID signal was maintained in both (Fig. 1C). Thus, this approach has generated the first clean null alleles in the *GluRIIA* and *GluRIIB* receptor subunits.

Next, we used immunostaining and electrophysiology to determine the composition and functionality of receptive fields exclusively composed of GluRA or GluRB. In *GluRIIA* mutants, a compensatory ∼170% enhancement in GluRIIB intensity was found, with no significant change in the common GluRIID subunit (Fig. 1C,D). However, mEPSC amplitude, which reflects the postsynaptic current induced by the spontaneous release of single synaptic vesicles, was reduced by over 50% without changing mEPSC frequency (Fig. 1E,F), as expected by exclusive expression of the rapidly desensitizing GluRB receptors and as observed in previous *GluRIIA* mutant studies (Diantonio et al., 1999; Petersen et al., 1997). Conversely, GluRIIA levels were similarly increased in *GluRIIB* mutants, with no overall change in GluRIID (Fig. 1C,D). This increased GluRA expression was reflected by a large increase in mEPSC amplitude and charge transfer (Fig. 1E,F and Table S1). Taken together, these experiments suggest that GluRA and GluRB receptors compete to establish postsynaptic receptive fields without changing total GluR abundance at postsynaptic compartments.

### Excess glutamate release adaptively reduces GluR abundance

GluRA and GluRB receptors compete to establish postsynaptic receptive fields, but it is not clear what physiologic signals regulate this competition. We hypothesized that the levels of synaptically released glutamate may modulate postsynaptic GluR abundance and/or composition. We first eliminated all synaptic glutamate release to determine whether GluR abundance or organization requires synaptic activity or glutamate itself. To accomplish this, we used a botulinum neurotoxin (BoNT) transgene that targets the SNARE protein Syntaxin for cleavage at release sites to block all neurotransmitter release (Han et al., 2022). Specifically, selective expression of BoNT-C in a subset of motor neurons using the driver OK319-GAL4 eliminates all miniature and evoked all glutamate release without impacting muscle target innervation or synaptic growth (Fig. 2A,B; (Han et al., 2022)). We then quantified GluRIIA, GluRIIB, and GluRIID immuno-intensity levels and found no significant difference in their levels or organization compared to wild type controls (Fig. 2C-E). Thus, glutamate release from presynaptic release sites is not required to establish or maintain GluRA or GluRB abundance at the *Drosophila* NMJ.

**Figure 2:**
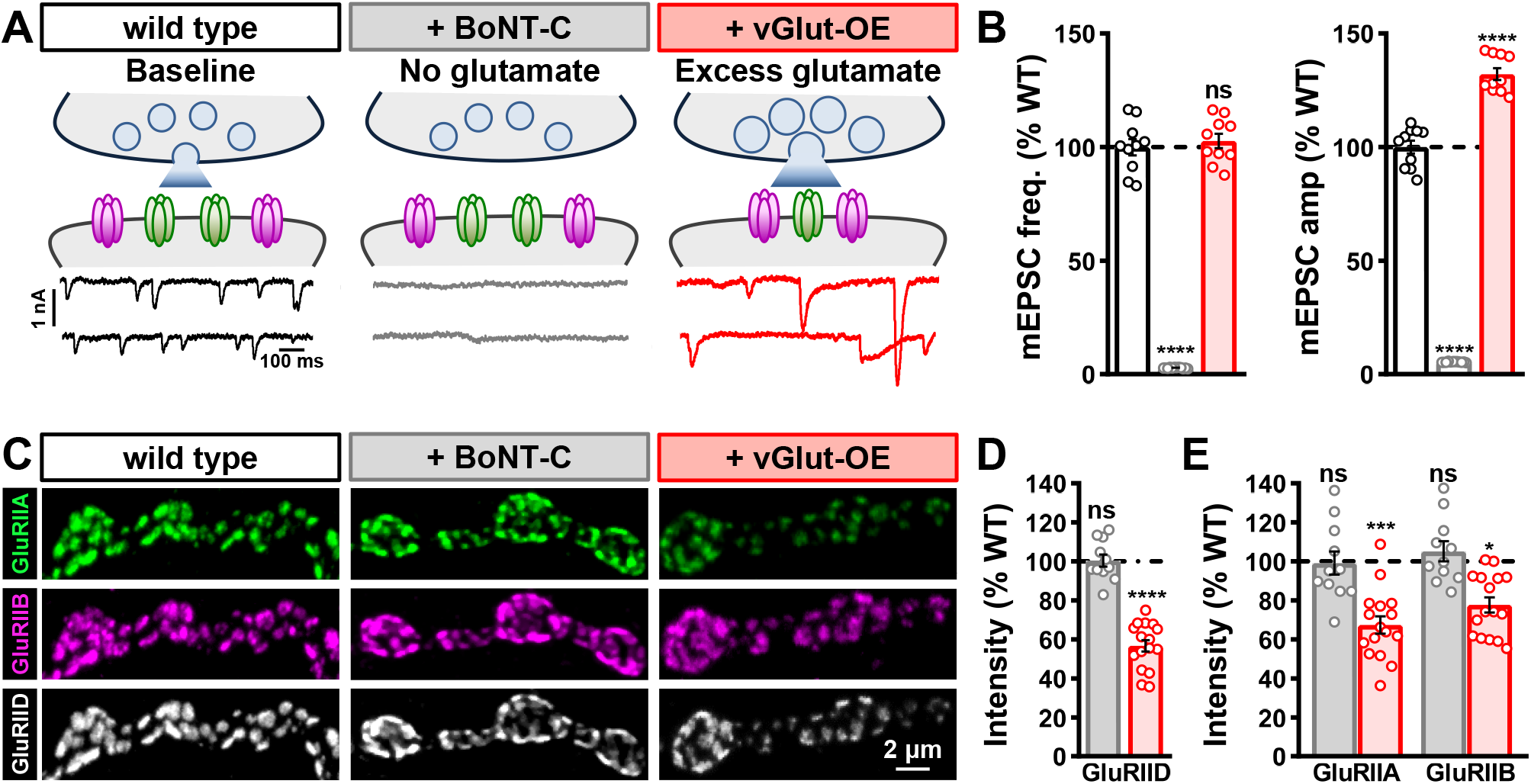
Excess presynaptic glutamate release induces a compensatory reduction in postsynaptic GluR abundance. **(A)** Schematics and representative mEPSC traces of wild type, NMJs with no synaptic glutamate release due to *BoNT-C* expression (*w*;*OK319-GAL4*/*+*;*UAS-BoNT-C*/*+*), and NMJs releasing excess glutamate due to *vGlut* overexpression in motor neurons (vGlut-OE: *w;OK371-Gal4/UAS-vGlut*). **(B)** Quantification of mEPSC frequency and amplitude in the indicated genotypes. Note that while BoNT-C eliminates all synaptic vesicle release and miniature activity, vGlut-OE leads to enhanced quantal size, as expected. **(C)** Representative images of wild type, BoNT-C, and vGlut-OE NMJs immunostained with anti-GluRIIA, -GluRIIB, and -GluRIID. While GluR abundance is unchanged in the absence of glutamate release, excess glutamate induces a compensatory reduction in GluR abundance. **(D)** Quantification of mean fluorescence intensity of individual GluRIID puncta in the indicated genotypes normalized to wild type values, indicating a reduction in total GluR abundance in vGlut-OE. **(E)** Quantification of anti-GluRIIA and -GluRIIB mean fluorescence intensity. Note that significant reductions in both GluRA and GluRB receptors are observed in vGlut-OE NMJs.

Next, we tested whether enhanced presynaptic glutamate release impacts postsynaptic GluR fields. Neuronal overexpression of the *vesicular glutamate transporter* (*vGlut*) increases the size of synaptic vesicles and leads to a concomitant increase in the abundance of glutamate emitted from single synaptic vesicles (Daniels et al., 2004; Li et al., 2018, 2021). Transgenic overexpression of *vGlut* in motor neurons using OK371-GAL4 (vGlut-OE) enhanced mEPSC amplitude, as expected (Fig. 2A,B). Interestingly, total GluR abundance, as assessed by GluRIID immunofluorescence intensity, was reduced by ∼50% in vGlut-OE (Fig. 2C,D), with both GluRIIA and GluRIIB levels significantly diminished at postsynaptic compartments (Fig. 2C,E). Thus, excess presynaptic glutamate release induces a compensatory reduction in postsynaptic GluR abundance.

### GluRA receptors are homeostatically regulated in the absence of GluRB competition

Although excess glutamate appears to adaptively reduce postsynaptic GluR levels, this apparent change in GluR abundance is compensatory but not homeostatic. In particular, mEPSC amplitudes are still increased in vGlut-OE, indicating that the reduction in GluRA and GluRB is not sufficient to maintain baseline miniature activity. We noticed that vGlut-OE reduces GluRA levels more than GluRB, and hypothesized that GluRA vs GluRB competition may obscure the individual responses of GluRA vs GluRB to excess glutamate. Therefore, we next sought to isolate the behavior of GluRA receptors to excess glutamate in the absence of GluRB receptors and vice versa.

To characterize the behavior of GluRA or GluRB receptors in the absence of subtype competition, we manipulated glutamate in *GluRIIA* or *GluRIIB* null mutants, where NMJs were composed exclusively of GluRB or GluRA. First, we found that *GluRIIA*- or *GluRIIB*-mutant synapses devoid of glutamate release by BoNT-C expression had no significant impact on GluR levels in GluRB- or GluRA-only NMJs (Table S1). Next, we examined the impact of vGlut-OE on *GluRIIA* mutant NMJs, which exclusively express GluRB receptors (Fig. 3A,B). mEPSCs were reduced by ∼50% in *GluRIIA* mutants alone compared to wild type, as expected, but were increased by over 70% in *GluRIIA*+vGlut-OE compared to baseline values (*GluRIIA* mutants; Fig. 3A). Correspondingly, immunostaining revealed that GluRIIB and GluRIID levels did not change in *GluRIIA*+vGlut-OE compared to *GluRIIA* mutants alone (Fig. 3B). These data indicate that in the absence of GluRA receptors, GluRB receptors become insensitive to and are not adaptively modulated by excess glutamate release.

**Figure 3:**
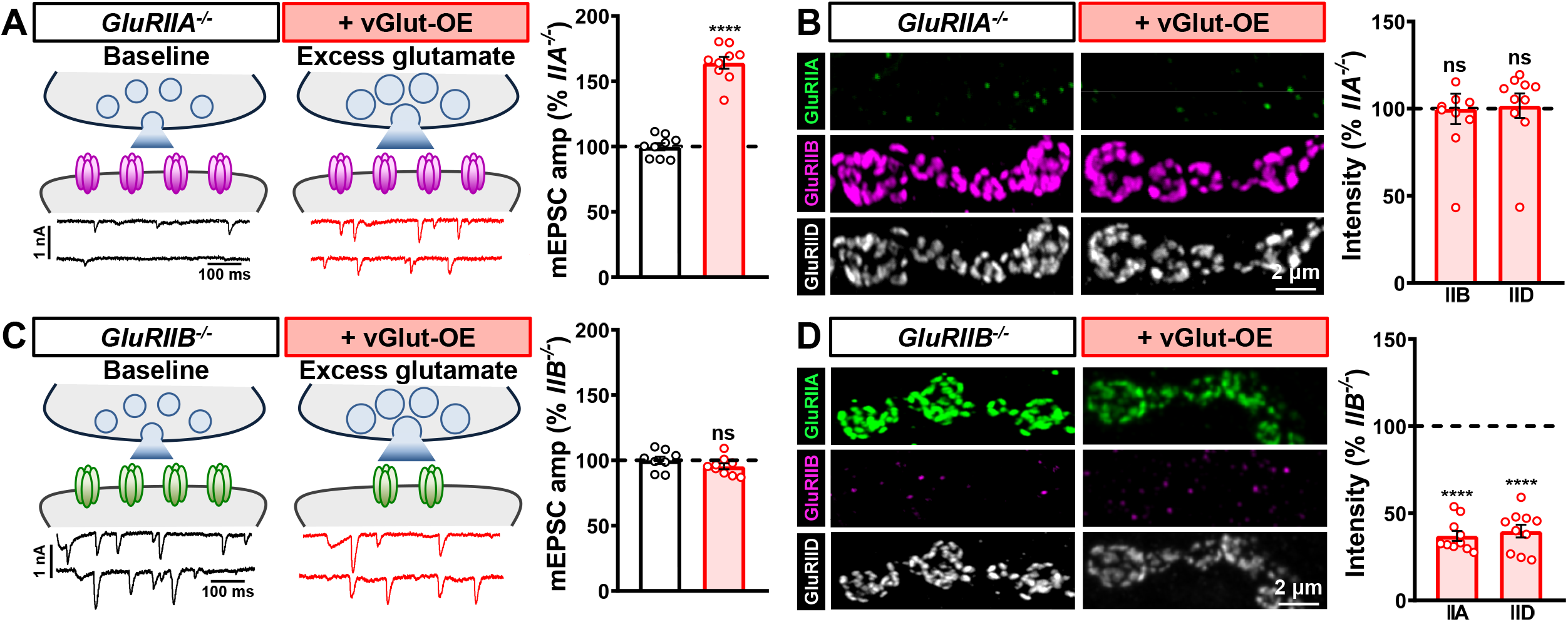
Excess glutamate selectively and homeostatically downregulates GluRA in the absence of GluRB. **(A)** Schematics and representative mEPSC traces of NMJs containing only GluRB receptors (*GluRIIA^-/-^* mutants: *w;GluRIIA^PV3^*) at baseline and following vGlut-OE (*GluRIIA*+vGlut-OE: *w;GluRIIA^PV3^,UAS-vGlut/GluRIIA^PV3^,OK371-Gal4*). Right: Quantification of mEPSC amplitude in both genotypes normalized to baseline (*GluRIIA* mutants). Note that mEPSC amplitude is significantly enhanced in *GluRIIA*+vGlut-OE over *GluRIIA* mutants alone. **(B)** Representative images of GluRs immunostained in the indicated genotypes. Right: Quantification of GluR mean fluorescence intensity in *GluRIIA*+vGlut-OE normalized to *GluRIIA* mutants alone. In the absence of GluRA receptors, excess glutamate release does not significantly change GluRB abundance. **(C)** Schematics and representative mEPSC traces of NMJs containing only GluRA receptors (*GluRIIB^-/-^* mutants: *w;GluRIIB^SP14^*) at baseline and following vGlut-OE (*GluRIIB*+vGlut-OE: *w;GluRIIB^SP14^,UAS-vGlut/GluRIIB^SP14^,OK371-Gal4*). Right: Quantification of mEPSC amplitude in both genotypes normalized to baseline (*GluRIIB* mutants). Note that in this case, no significant change in mEPSC amplitude is observed despite excess glutamate released by vGlut-OE. **(D)** Representative images of GluRs immunostained in the indicated genotypes. Right: Quantification of GluR mean fluorescence intensity in *GluRIIB*+vGlut-OE normalized to *GluRIIB* mutants alone. In this situation, excess glutamate homeostatically downregulates GluRA abundance in the absence of competition with GluRB receptors.

Finally, we characterized the impact of excess glutamate in *GluRIIB* mutants. In these mutants, baseline mEPSC amplitudes were elevated by ∼50% compared to wild type due to the exclusive and enhanced expression of GluRA receptors (Fig. 3C,D). Remarkably, however, no significant difference in mEPSC amplitude was observed in *GluRIIB* mutants with excess glutamate release compared to *GluRIIB* mutants alone (Fig. 3C). This suggests that in the absence of GluRB, GluRA receptors are homeostatically downregulated by excess glutamate to maintain baseline miniature amplitude. Indeed, vGlut-OE led to a >60% reduction in GluRIIA levels compared to baseline (*GluRIIB* mutants; Fig. 3D). This quantitative reduction in GluRA abundance was sufficient in amplitude to explain the stable mEPSC values despite excess glutamate driven by vGlut-OE. Thus, the elimination of competition between GluRA and GluRB receptors reveals two distinct, subtype-specific responses to excess synaptic glutamate release at fly NMJ receptive fields: 1) GluRB receptors are immutable, remaining fixed and unresponsive to glutamate, while 2) GluRA receptors are plastic, precisely downregulated in response to excess synaptic glutamate to homeostatically maintain stable miniature activity.

### Ca^2+^ permeability through GluRA is necessary for homeostatic receptor scaling

Postsynaptic GluRA receptor abundance is homeostatically diminished by enhanced glutamate release. Ca^2+^ influx through GluRs and related signaling in postsynaptic compartments orchestrate a number of forms of plasticity that ultimately modulate GluR abundance and composition at postsynaptic receptive fields (Bayer and Schulman, 2019; Herring and Nicoll, 2016). GluRA receptors are Ca^2+^ permeable and pass the majority of glutamate-elicited currents at postsynaptic compartments of the *Drosophila* NMJ (Diantonio et al., 1999; Han et al., 2015). We therefore hypothesized that Ca^2+^ influx through GluRA receptors may be necessary for homeostatic receptor scaling induced by excess glutamate release.

To test this hypothesis, we generated a *GluRIIA* allele designed to specifically eliminate Ca^2+^ influx through GluRA using CRISPR/Cas9 gene editing. Like most Ca^2+^ permeable GluRs, the *GluRIIA* subunit encodes a neutral glutamine amino acid (Q) in the pore-forming M2 loop (Q615; Fig. 4A). To render GluRA Ca^2+^ impermeable while remaining permeable to other ions, we used CRISPR/Cas9 genome editing to mutate this glutamine to the positively charged amino acid arginine (R) at the endogenous *GluRIIA* locus to make *GluRIIA^Q615R^* alleles. In heterologous systems, this Q to R transition renders both AMPA- and Kainate-GluRs impermeable to Ca^2+^ (Hume et al., 1991; Köhler et al., 1993; Li et al., 2016; Ni, 2021). *GluRIIA^Q615R^* mutants are homozygous viable and healthy, exhibit normal synaptic transmission (miniature and evoked release), and express GluRA and GluRB receptors that do not significantly change in abundance compared to wild type (Fig. S1A-D), as expected. We then performed Ca^2+^ imaging at postsynaptic NMJ compartments using GCaMP8f targeted to postsynaptic densities (Fig. S2A, SynapGCaMP8f; (Han et al., 2022)) in wild type, *GluRIIA^PV3^*, *GluRIIA^Q615R^*, and *GluRIIB^SP14^* mutants, as well as vGlut-OE. Quantal Ca^2+^ events were reduced by ∼50% in *GluRIIA^PV3^* null mutants compared to wild type, and enhanced by ∼50% in *GluRIIB^SP14^* and vGlut-OE, as expected (Fig. S2B,C). Importantly, a ∼50% reduction in postsynaptic Ca^2+^ signals were observed in *GluRIIA^Q615R^* mutants, statistically indistinguishable from *GluRIIA* null mutants (Fig. S2B,C). Thus, *GluRIIA^Q615R^* mutants render GluRA receptors Ca^2+^ impermeable without impacting basal synaptic transmission or relative GluR composition or abundance.

**Figure 4:**
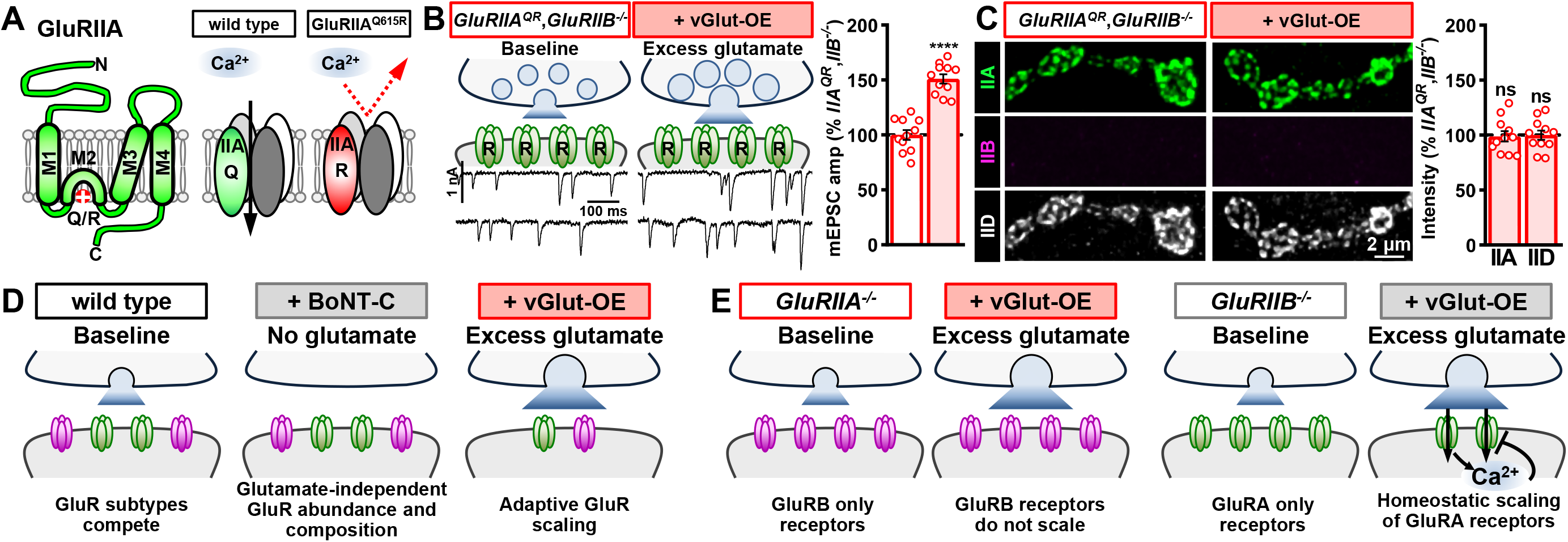
Homeostatic control of GluRA abundance by excess glutamate requires Ca^2+^ permeability. **(A)** Schematics illustrating GluRIIA subunit topology with the Q615R mutation engineered in the pore forming M2 domain, rendering GluRA receptors Ca^2+^ impermeable. **(B)** Schematics and representative mEPSC traces of NMJs containing only Ca^2+^ impermeable GluRA receptors (*GluRIIA^Q615R^,GluRIIB*: w;*GluRIIA^Q615R^*,*GluRIIB^SP5^*/*GluRIIA^Q615R^*,*GluRIIB^SP5^*;+) at baseline and following vGlut-OE (*GluRIIA^QR^*,*GluRIIB*+vGlut-OE: w;*OK371-GAL4*,*GluRIIA^Q615R^*,*GluRIIB^SP5^*/*UAS-vGlut*,*GluRIIA^Q615R^*,*GluRIIB^SP5^*;*+*). Right: Quantification of mEPSC amplitude in both genotypes normalized to baseline (*GluRIIA^Q615R^,GluRIIB* mutants). Note that a significant increase in mEPSC amplitude is observed following excess glutamate released by vGlut-OE, which was not observed with Ca^2+^ permeable GluRA (Fig. 3C). **(C)** Representative images of GluRs immunostained in the indicated genotypes. Right: Quantification of GluR mean fluorescence intensity in *GluRIIA^QR^*,*GluRIIB*+vGlut-OE normalized to *GluRIIA^QR^*,*GluRIIB* mutants alone. Without Ca^2+^ permeability, excess glutamate no longer downregulates GluRA abundance in the absence of competition with GluRB receptors. **(D)** Summary of glutamate loss or enhancement on GluR receptive fields when GluRA and GluRB are in competition. **(E)** Summary of the impact of excess glutamate on GluR receptive fields when GluR subtype competition is lost.

Finally, we determined whether homeostatic GluRA receptor scaling is affected when GluRA receptors are Ca^2+^ impermeable. First, we found that in *GluRIIA^Q615R^* mutants, GluRA and GluRB receptors behave like wild type at baseline and after vGlut-OE, with reductions in both GluRA and GluRB (Fig. S1E-H). To specifically examine homeostatic GluRA scaling, we generated CRISPR-induced *GluRIIB* null mutations in *GluRIIA^Q615R^* mutant backgrounds, leading to synapses composed entirely of Ca^2+^-impermeable GluRA receptors (Fig. 4B). In *GluRIIA^Q615R^*,*GluRIIB* double mutants, baseline mEPSC amplitudes were enhanced, as expected, but were now further enhanced by ∼50% when combined with vGlut-OE (Fig. 4B). Consistently, we observed no change in GluRIIA or GluRIID levels in *GluRIIA^Q615R^*, *GluRIIB* double mutants with vGlut-OE compared to *GluRIIA^Q615R^* mutants alone (Fig. 4C). These data indicate that when GluRA is rendered impermeable to Ca^2+^ and competition with GluRB is eliminated, excess glutamate is no longer capable of inducing homeostatic reductions in GluRA levels at postsynaptic compartments. We summarize these results in Fig. 4D,E.

## DISCUSSION

By generating null mutations in *GluRIIA* and *GluRIIB* subunits, we have shown a competition exists between GluR subtypes that establishes stable postsynaptic fields. While synaptically released glutamate is not required to organize this process, excess glutamate triggers an adaptive downscaling of both GluR subtypes in the postsynaptic compartment. However, when this GluR-subtype competition is eliminated, a clear and distinctive relationship is revealed between excess glutamate and GluR plasticity: GluRB receptors become completely insensitive to excess glutamate, with stable GluRB levels maintained. In contrast, GluRA receptors constitute the “plastic” receptor subtype, homeostatically tuned to excess synaptic glutamate release to maintain stable miniature activity. Further, Ca^2+^ influx through GluRA receptors is a key transducer of this signaling system, rendering GluRA non-plastic when Ca^2+^ permeability is lost. Together, these results highlight the interplay between GluR subtype competition, synaptic glutamate, and Ca^2+^ signaling at postsynaptic compartments and reveals the existence of homeostatic receptor scaling at the *Drosophila* NMJ, a model glutamatergic synapse.

Several layers of regulation operate at postsynaptic compartments to establish GluR receptor fields at the fly NMJ. First, the relative level of GluR transcription and translation between subtypes can ultimately set GluRs at synapses. This is demonstrated by overexpression of either *GluRIIA* or *GluRIIB* subunits, which can saturate the entire GluR field at postsynaptic compartments and lead to the concomitant loss of the other GluR subtype (Diantonio et al., 1999; Li et al., 2018; Marrus et al., 2004). Second, post-translational processes, mediated by such factors as enzymatic cleavage, phosphorylation, and degradation modulate GluR activity and abundance. For example, Ca^2+^-dependent protein cleavage by Calpain, phosphorylation control by p21-activated kinase (Pak), and proteosomal degradation by the E3 ubiquitin ligase adapter Diablo have all been shown to modulate GluRs at the fly NMJ (Metwally et al., 2019; Wang et al., 2016), which parallel findings in vertebrates (Hawasli et al., 2007; Murata and Constantine-Paton, 2013; Simpkins et al., 2003; Verhagen et al., 2000).

Third, while glutamate released from synaptic vesicles is not necessary to establish or maintain GluR fields in *Drosophila* (Fig. 2) or in mammals (Sando et al., 2017; Sigler et al., 2017), there is evidence that ambient glutamate modulation from non-vesicular glutamate release and glial transporters might regulate GluR clustering and receptor field size (Augustin et al., 2007; Featherstone et al., 2002), while excess vesicular glutamate release triggers adaptive reductions in GluRs (Fig. 2). There is also evidence that correlated or diminished activity selectively regulate GluRA abundance (Ljaschenko et al., 2013). It will be of interest to determine how these many layers of control intersect and are coordinated to establish GluR fields during development and remodel in plasticity.

There appears to be a hierarchy of regulatory steps controlling GluR plasticity in response to excess glutamate. While excess glutamate downregulates both GluRA and GluRB abundance when both receptor subtypes are present, this plasticity is adaptive but not homeostatic – miniature amplitude is still enhanced. However, when GluRA vs GluRB competition is eliminated, a complete distinction in GluR behavior is revealed: GluRB is no longer responsive to excess glutamate, while GluRA receptors are sensitively tuned to glutamate to now homeostatic control miniature activity. Ca^2+^ influx through GluRA appears crucial to this plasticity, where loss of this secondary messenger converts GluRA to behave like static GluRB receptors. It is possible that excess glutamate drives Ca^2+^-related signaling through GluRA in postsynaptic compartments that ultimately acts on both GluRA and GluRB when both receptor subtypes are present, and this signaling may be lost in GluRB-only NMJs. Although the downstream effectors that respond to excess glutamate and Ca^2+^ to modulate postsynaptic GluRs are not known, an attractive candidate is the auxiliary KAR subunit Neto, which regulates GluR abundance at the fly NMJ (Kim and Serpe, 2013; Kim et al., 2012). Interestingly, KARs in mammals are also under homeostatic control (Yan et al., 2013), where the auxiliary subunit Neto controls key properties of these receptors (Tomita et al., 2012).

What purpose might two GluR subtypes, differing in their current amplitudes, biophysics, and plasticity subserve? One idea is that GluRB receptors provide a basal signal at postsynaptic compartments to maintain synaptic dialogue, while GluRA receptors are the potent subtype that sets synaptic strength, drives muscle contractions, and is targeted for plasticity. These differential functions may be reflected in their distinct subsynaptic localizations, with GluRA enriched opposite active zone centers where glutamate is released, while GluRB is enriched in the outside periphery of these areas (Han et al., 2022; Marrus et al., 2004; Muttathukunnel et al., 2021). Another possibility, not mutually exclusive, is that GluRBs serve as ‘back-up’ receptors to maintain NMJ transmission and locomotion when GluRAs are blocked, a phenomenon that occurs naturally at larval NMJs due to toxins injected by parasitoid wasps and other organisms (Eldefrawi et al., 1988; Hwang et al., 2007; Karst and Piek, 1991). Indeed, presynaptic homeostatic potentiation, a conserved form of retrograde plasticity modeled at the fly NMJ, is induced when GluRA is lost or pharmacologically inhibited to maintain stable NMJ excitation (Goel and Dickman, 2021). Hence, stable GluRB receptors provide robustness to buffer NMJ function from perturbations while also allowing flexibility for GluRA receptors to dynamically change with plasticity.

Although the *Drosophila* NMJ has long been used as a model to study presynaptic forms of adaptive plasticity such homeostatic potentiation and depression (Frank et al., 2020; Goel and Dickman, 2021), recent work has found parallel modes of adaptive plasticity that target postsynaptic GluR abundance at this model glutamatergic synapse. In addition to excess glutamate targeting GluRs, at least three additional examples of adaptive GluR plasticity have been observed at the fly NMJ. First, in synaptic undergrowth mutants, where presynaptic innervation is reduced, overall synaptic strength can be maintained at least in some cases through an adaptive enhancement of postsynaptic GluR abundance (Goel et al., 2019). Second, when innervation by a single motor neuron is biased at adjacent muscles, stable synaptic strength is maintained through both pre- and post-synaptic mechanisms (Davis and Goodman, 1998), with a homeostatic increase in postsynaptic GluRs necessary at hypo-innervated NMJs (Goel et al., 2020). Third, activation of injury-related signaling in motor neurons induces a downregulation in postsynaptic GluR abundance to adaptively reduce the set point of synaptic strength (Goel and Dickman, 2018). These examples suggest an unanticipated level of postsynaptic plasticity exists at the *Drosophila* NMJ which, when combined with the sophisticated genetic and functional tools available, highlights the great potential for the *Drosophila* NMJ to illuminate how GluR subtype competition and presynaptic function target GluRs for adaptive modulation.

## MATERIALS AND METHODS

### Fly stocks

*Drosophila* stocks were raised at 25°C on standard molasses food. The *w^1118^* strain is used as the wild type control unless otherwise noted as this is the genetic background in which all genotypes are bred. Details of all fly stocks used, including their sources, are listed in Table S2.

### Molecular Biology

*GluRIIA^PV3^, GluRIIA^PV7^, GluRIIB^SP5^*, and *GluRIIB^SP14^* mutants were generated using a CRISPR/Cas9 genome editing strategy as described (Kikuma et al., 2017). For *GluRIIA* mutants, a TKO stock was obtained from BDSC (#68059) that ubiquitously expressed a sgRNA (5’ CAATCGCACCGACGTAATGTTGG 3’) targeting the sixth exon of the *GluRIIA* locus (Fig. 1B). To generate *GluRIIB* mutants, we generated two independent sgRNA lines that targeted the first and sixth exons (sgRNA1: 5’ GGTGTCTTCATTGGCGCCGCTGG 3’; sgRNA2: 5’ CATTGATGGATTCTACTCCCGGG 3’) and cloned each into the pU63 vector (#49410; Addgene). Constructs were sent to BestGene Inc. (Chino Hill, CA) for targeted insertion into the VK18 attP site on the second chromosome. sgRNA flies were crossed to a *nos-Cas9* line (#54591;BDSC) on the second chromosome to induce active germline CRISPR mutagenesis, and 20 independent lines generated from each sgRNA were screened by PCR for mutations. This identified at least 8 independent indel mutations for each sgRNA that shifted the open reading frame, with *GluRIIA^PV3^, GluRIIA^PV7^, GluRIIB^SP5^*, and *GluRIIB^SP14^* alleles kept for additional analysis (Fig. 1B). An identical strategy was used to generate *GluRIIB* mutations in the *GluRIIA^Q615R^* background.

To generate the Ca^2+^ impermeable *GluRIIA^Q615R^* allele, a sequence containing 1 kb homology arms flanking the *GluRIIA* genomic region with the Q615R point mutation was inserted into pHD-DsRed vector (#51434; Αddgene) as the CRISPR donor. Two single guide RNAs (gRNA1: gaacaactcgacttggctga, gRNA2: ggtgggctccatcatgcaac) were inserted together into the pAC-U63-tgRNA (#112811; Addgene) vector with intervening tRNA(F+E) sequences for expressing multiple gRNAs (Poe et al., 2019). The donor construct and the gRNA construct were then co-injected into a *nos-Cas9* (#78782; BDSC) fly strain by Well Genetics (Taipei City, Taiwan (R.O.C.)) to generate the *GluRIIA^Q615R^* mutant by homology-directed repair. Successful CRISPR fly lines were selected by P3>DsRed expression in eyes and confirmed by PCR. DsRed with flanking PBac sequence was then removed by PBac-mediated excision using the *Tub>PBac* fly strain (#8283, BDSC).

### Electrophysiology

All dissections and two-electrode voltage clamp (TEVC) recordings were performed as described (Kikuma et al., 2019)using modified hemolymph-like saline (HL-3) containing: 70mM NaCl, 5mM KCl, 10mM MgCl_2_, 10mM NaHCO_3_, 115mM Sucrose, 5mM Trehelose, 5mM HEPES, and 0.5mM CaCl_2_, pH 7.2, from cells with an initial resting potential between −60 and −75 mV, and input resistances >6 MΩ. Recordings were performed on an Olympus BX61 WI microscope using a 40x/0.80 NA water-dipping objective and acquired using an Axoclamp 900A amplifier, Digidata 1440A acquisition system and pClamp 10.5 software (Molecular Devices). Miniature excitatory postsynaptic currents (mEPSCs) were recorded in the absence of any stimulation with a voltage clamp of −80 mV, and low pass filtered at 1 kHz. All recordings were made on abdominal muscle 6, segments A2 or A3 of third-instar larvae with the leak current never exceeding 5 nA. mEPSCs were recorded for 60 seconds and analyzed using MiniAnalysis (Synaptosoft) and Excel (Microsoft) software. The average mEPSC amplitude and total charge transfer values for each NMJ were obtained from approximately 100 events in each recording.

### Immunocytochemistry

Third-instar larvae were dissected in ice cold 0 Ca^2+^ HL-3 and immunostained as described (Chen and Dickman, 2017; Goel et al., 2017; Li et al., 2021). In brief, larvae were either fixed in Bouin’s fixative for 5 min (Sigma, HT10132-1L), 100% ice-cold ethanol for 5 min, or 4% paraformaldehyde (PFA) for 10 min. Larvae were then washed with PBS containing 0.1% Triton X-100 (PBST) for 30 min, blocked with 5% Normal Donkey Serum followed by overnight incubation in primary antibodies at 4°C. Preparations were then washed 3x in PBST, incubated in secondary antibodies for 2 hours, washed 3x in PBST, and equilibrated in 70% glycerol. Prior to imaging, samples were mounted in VectaShield (Vector Laboratories). Details of all antibodies, their source, dilution used, and references are listed in Table S2.

### Imaging and analysis

Samples were imaged using a Nikon A1R Resonant Scanning Confocal microscope equipped with NIS Elements software and a 100x APO 1.4NA oil immersion objective using separate channels with four laser lines (405 nm, 488 nm, 561 nm, and 647 nm) as described (Perry et al., 2017). For fluorescence intensity quantifications of GluRIIA, GluRIIB and GluRIID, z-stacks were obtained on the same day using identical gain and laser power settings with z-axis spacing between 0.15-0.20 µm for all genotypes within an individual experiment. Maximum intensity projections were utilized for quantitative image analysis using the general analysis toolkit of NIS Elements software. Immunofluorescence intensity levels were quantified by applying intensity thresholds and filters to binary layers in the 405 nm, 488 nm, and 561 nm channels. The mean intensity for each channel was quantified by obtaining the average total fluorescence signal for each individual punctum and dividing this value by the puncta area. A mask was created around the HRP channel, used to define the neuronal membrane, and only puncta within this mask were analyzed to eliminate background signals. All measurements based on confocal images were taken from NMJs acquired from at least six different animals.

### Ca^2+^ imaging and analysis

Third-instar larvae were dissected in ice-cold saline. Imaging was performed in modified HL-3 saline with 1.5 mM Ca^2+^ added using a Zeiss Examiner A1 widefield microscope equipped with a 63x/1.0 NA water immersion objective as described (Han et al., 2022). NMJs on muscle 6 were imaged at a frequency of 100 fps (512 x 256 pixels) with a 470 nm LED light source (Thorlabs) using a PCO sCMOS4.2 camera. Spontaneous Ca^2+^ events were imaged at NMJs during 120 sec imaging sessions from at least two different larvae.

Evoked Ca^2+^ events were induced by delivering 10 electrical stimulations at 0.5 Hz. Horizontal drifting was corrected using ImageJ plugins (Kang Li, 2008) and imaging data with severe muscle movements were rejected as described (Ding et al., 2019). Three ROIs were manually selected using the outer edge of terminal Ib boutons observed by baseline GCaMP signals with ImageJ (Rueden et al., 2017; Schindelin et al., 2012). Ib and Is boutons were defined by baseline GCaMP8f fluorescence levels, which are 2-3 fold higher at Ib NMJs compared to their Is counterparts at a particular muscle. Fluorescence intensities were measured as the mean intensity of all pixels in each individual ROI. ΔF for a spontaneous event was calculated by subtracting the baseline GCaMP fluorescence level F from the peak intensity of the GCaMP signal during each spontaneous event at a particular bouton as previously detailed. ΔF/F was calculated for each spontaneous Ca^2+^ transient event as detailed (Han et al., 2022).

### Statistical analysis

Data were analyzed using GraphPad Prism (version 7.0), MiniAnalysis (Synaptosoft), or Microsoft Excel software (version 16.22). Sample values were tested for normality using the D’Agostino & Pearson omnibus normality test which determined that the assumption of normality of the sample distribution was not violated. Data were then compared using either a one-way ANOVA and tested for significance using a Tukey’s multiple comparison test or using an unpaired 2-tailed Student’s t-test with Welch’s correction. In all figures, error bars indicate ±SEM, with the following statistical significance: p<0.05 (*), p<0.01 (**), p<0.001 (***), p<0.0001 (****); ns=not significant. Additional statistics and sample number values (n) for all experiments are summarized in Table S1.

## ACKNOWLEDGEMENTS

We thank Aaron DiAntonio (Washington University, MO, USA) for sharing *Drosophila* stocks. We acknowledge the Developmental Studies Hybridoma Bank (Iowa, USA) for antibodies used in this study and the Bloomington Drosophila Stock Center for fly stocks (NIH P40OD018537). We thank Giwoo Kim, Veronica Haro-Acosta, Surbhi Trivedi, and Chun Chien for assistance in validating reagents. This work was supported by a grant from the National Institutes of Health (NS111414) to DD.

## AUTHOR CONTRIBUTIONS

PG and YH obtained all experimental data. NT contributed genetic experiments, SN and MS contributed technical support, and SP generated the initial *GluRIIB* mutant alleles. PG, YH, and DD analyzed and interpreted all data. The manuscript was written by DD with feedback from YH and PG.

---

**Figure S1:**
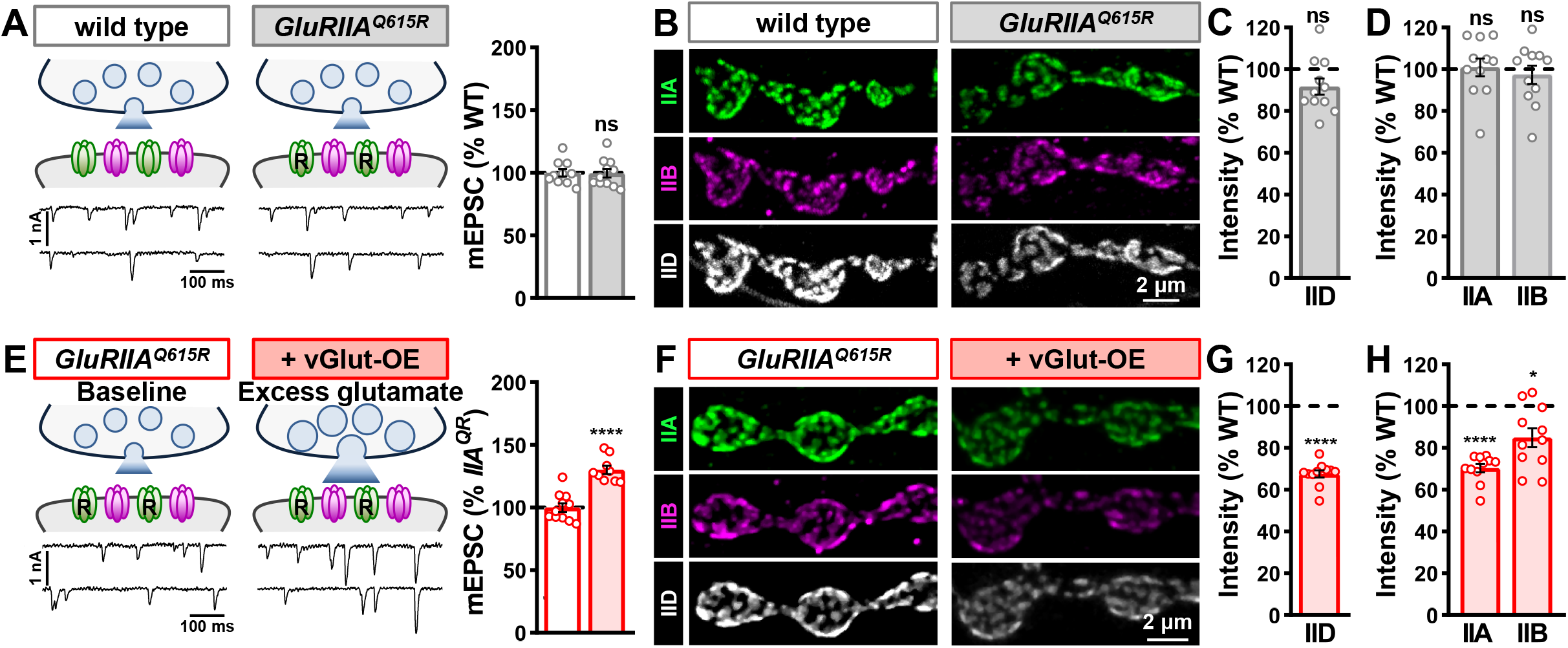
Ca^2+^ impermeable GluRA receptors behave like wild type at baseline and following *vGlut* overexpression. **(A)** Schematics and representative mEPSC traces of wild type and *GluRIIA^Q615R^* mutants. Right: Quantification of mEPSC amplitude in both genotypes normalized to wild type. No significant difference in mEPSC amplitude is observed. **(B)** Representative images of GluRs immunostained in the indicated genotypes. **(C,D)** Quantification of GluRIID (C), and GluRIIA and GluRIIB (D) mean fluorescence intensity in *GluRIIA^Q615R^* normalized to wild type. No significant difference in GluRA, GluRB, or total GluR intensity is found. **(E)** Schematics and representative mEPSC traces of *GluRIIA^Q615R^* at baseline and following *vGlut* overexpression (*GluRIIA^Q615R^*+vGlut-OE: *w;GluRIIA^Q615R^, OK371-Gal4/GluRIIA^Q615R^*, UAS-vGlut). Right: Quantification of mEPSC amplitude in both genotypes normalized to baseline (*GluRIIA^Q615R^* mutants). A similar increase in mEPSC amplitude is observed compared vGlut-OE at wild-type NMJs (Fig. 2). **(F-H)** Representative images (F) and quantification (G,H) of GluRs immunostained in the indicated genotypes. Quantification of GluR mean fluorescence intensity is normalized to wild type. GluRA, GluRB, and total GluR intensity is reduced to similar levels in *GluRIIA^Q615R^*+vGlut-OE as observed in wild type+vGlut-OE (Fig. 2).

**Figure S2:**
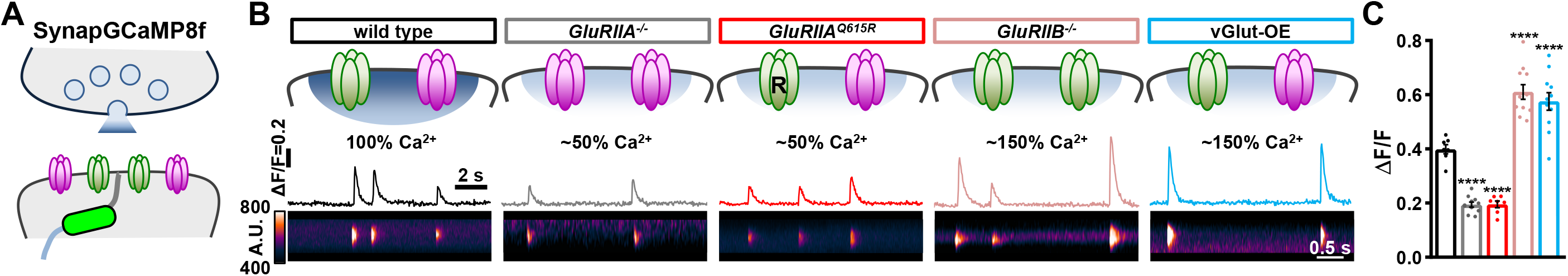
Engineering and validation of a Ca^2+^ impermeable *GluRIIA* allele. **(A)** Schematic illustrating the postsynaptic Ca^2+^ indicator SynapGCaMP8f localized to postsynaptic NMJ compartments. **(B)** Schematics and Ca^2+^ imaging traces of SynapGCaMP8f combined with the indicated genotypes. Line scans below are derived from postsynaptic GCaMP8f images of individual spontaneous Ca^2+^ transients. Note that while Ca^2+^ is reduced by ∼50% in *GluRIIA* null or Ca^2+^ impermeable *GluRIIA^Q615R^* mutants, Ca^2+^ levels are enhanced by ∼50% in *GluRIIB* null mutants or vGlut-OE. **(C)** Quantification of fluorescence intensity (ΔF/F) changes from spontaneous Ca^2+^ transient events at individual boutons in the indicated genotypes. Genotypes: wild type (*w*;*MHC-SynapGCaMP8f*/*+*), *GluRIIA^-/-^* (*w*;*GluRIIA^pv3^*;*MHC-SynaptoGCaMP8f*/*+*), *GluRIIA^Q615R^* (*w*;*GluRIIA^Q615R^*;*MHC-SynaptoGCaMP8f*/*+*), *GluRIIB^-/-^* mutants (*w*;*GluRIIB^SP5^*;*MHC-SynapGCaMP8f*/*+*), and vGlut-OE (*w*;*vGlut-Gal4*/*UAS-vGlut*;*MHC-SynapGCaMP8f*/*+*).

**Supplementary Table 1:**
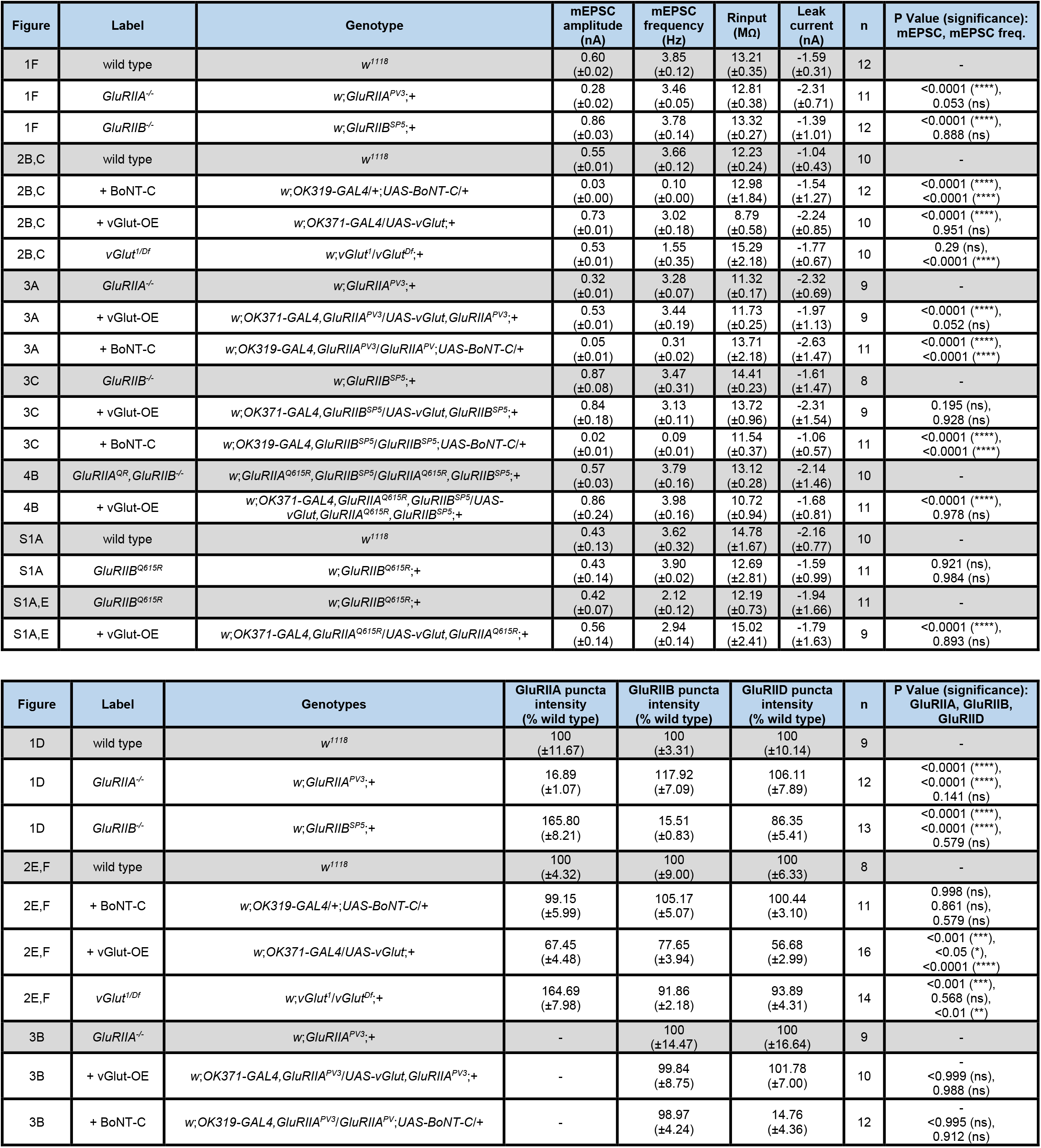
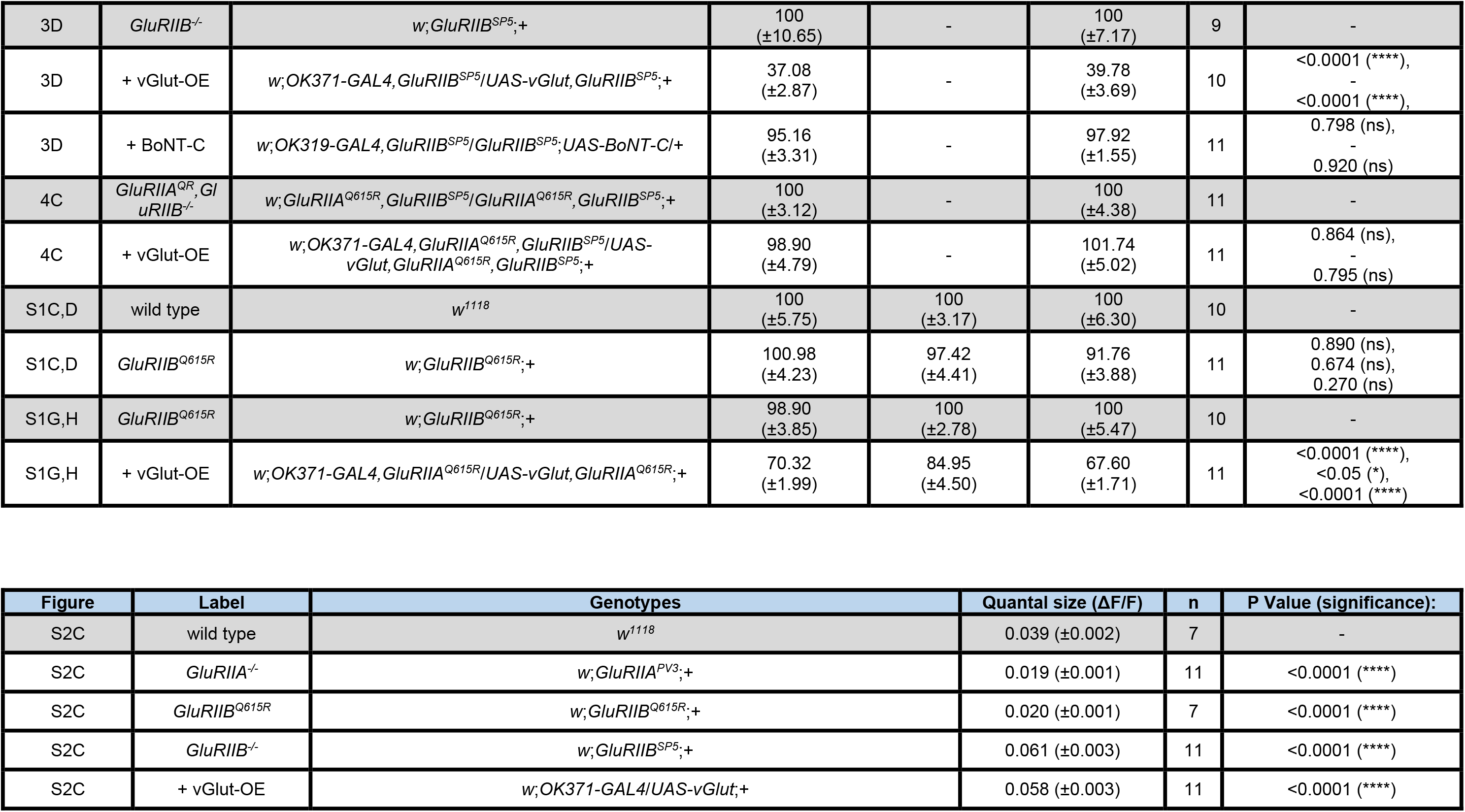
Absolute values for normalized data and additional statistical details. The figure and panel, genotype, and conditions are noted. Average values (with standard error of the mean noted in parentheses), data samples (n), and statistical significance tests and values are shown for all data.

## STAR★METHODS

### KEY RESOURCES TABLE

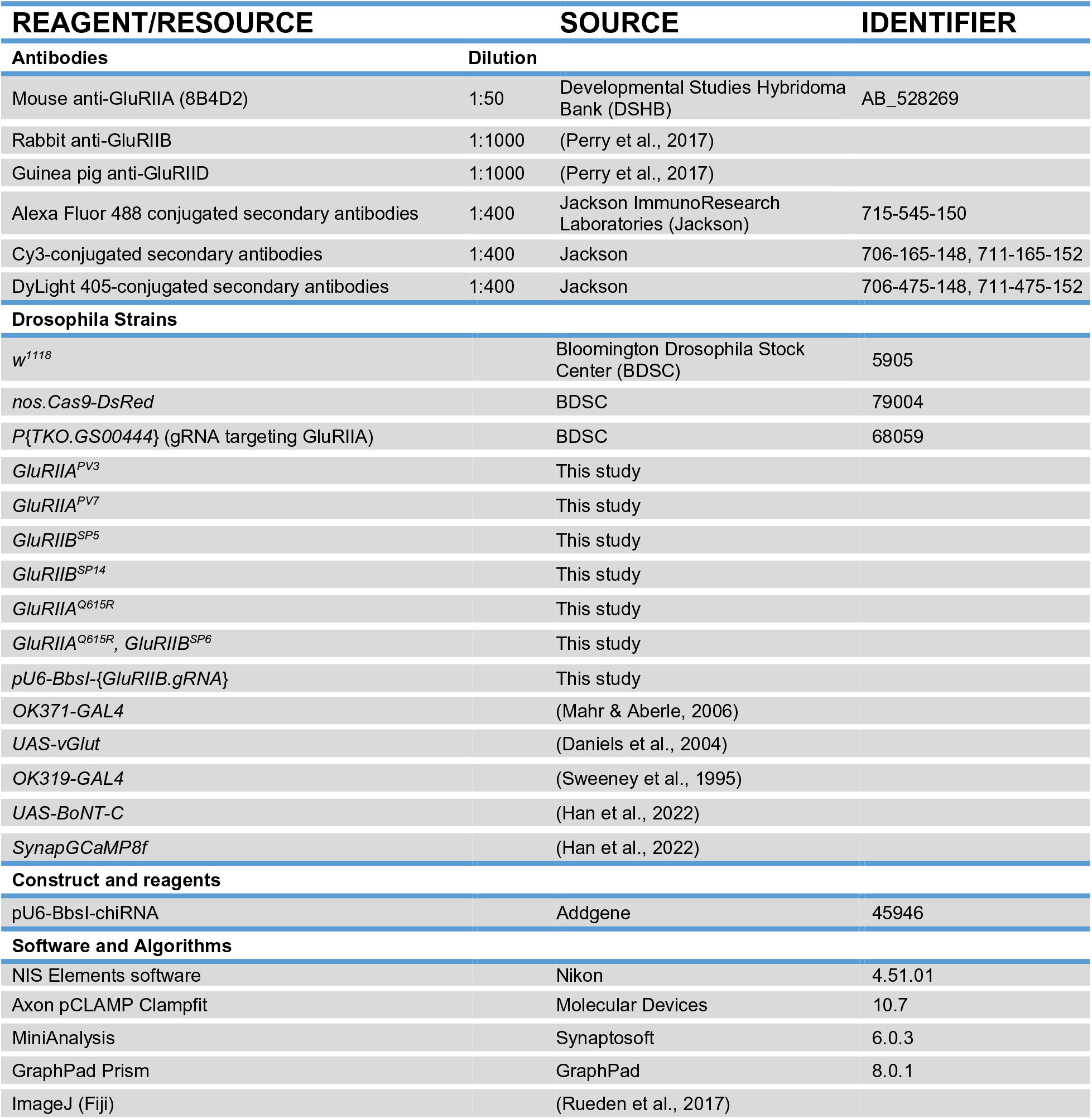

